# An extension of Modular Response Analysis for global perturbations and robust connectivity inference of gene regulatory networks

**DOI:** 10.64898/2026.06.02.729263

**Authors:** Gabriel Jimenez-Dominguez, Benjamin Audit, Pierre Borgnat, Patrice Ravel, Jean-Michel Arbona

## Abstract

Understanding how gene regulatory networks respond to global cell perturbations remains a central challenge in systems biology and network inference. Modular Response Analysis (MRA) provides a mathematical framework to infer gene-to-gene directed connectivity graphs from perturbation experiments; however, classical MRA captures direct gene-to-gene influences, and does not explicitly account for global stimuli that simultaneously change the graph. Here, we introduce MRA^+^, an extension of MRA, that incorporates the effect of global perturbations into gene-to-gene graph inference. MRA^+^ assumes a sequential experimental design in which targeted gene perturbations are followed by the application of a global stimulus, enabling the separation of connectivity changes from direct gene induction. The method estimates network connectivity under induced conditions and quantifies gene-specific induction strengths, which represent contributions to expression changes arising from mechanisms external to the inferred network. In the case of single-cell expression data, we present a bootstrap strategy to assess the robustness of inferred connectivity coefficients and propose a complementary criterion based on sign stability to interpret weak or non-significant estimates. Together, these developments provide a general framework for robust inference of gene connectivity graphs in the presence of global perturbations, applicable to diverse biological and experimental contexts.

## I. Introduction

Gene regulatory networks (GRNs) are commonly inferred from perturbation experiments, which provide causal information about directed interactions between molecular components. Modular Response Analysis (MRA) is a well-established mathematical framework to estimate directed connectivity from steady-state perturbation data [1], [2]. Over the past two decades, several extensions of MRA have been proposed to improve robustness and flexibility [3], including regression-based formulations to better handle noise, higher-order models to capture nonlinear effects, and the incorporation of prior information into network reconstruction [4], [5], or to take into account temporal data [6]. For GRNs, many methods of graph inference have been proposed over the year [7], yet MRA keeps its relevance as the framework comes from a dynamical models of the regulation.

Despite these advances, MRA and its variants remain designed to characterise a single biological condition. In contrast, interest is often in the characterization of a global perturbation of a reference biological condition—such as chemical induction, environmental changes, or signaling stimuli—that simultaneously affect multiple components of the system. Disentangling the effects of the targeted single-gene perturbations from those arising from global stimuli remains a methodological challenge for usual methods of network inference.

In parallel, recent developments in single-cell perturbation technologies, such as Perturb-seq [8], now enable the measurement of transcriptional responses to large numbers of targeted perturbations at single-cell resolution. These experiments provide unprecedented statistical power but also introduce substantial variability and sampling noise, making robust estimation of connectivity coefficients a critical issue. Resampling strategies such as bootstrap analysis therefore play an increasingly important role in assessing the stability and interpretability of inferred network parameters.

Here we introduce MRA^+^ an extension of MRA that incorporates global perturbations into the inference framework. We also provide a systematic strategy to evaluate the robustness of inferred coefficients using bootstrap resampling. The method separates connectivity-driven effects from global induction effects. For single-cell expression data, we introduce a criterion to interpret weak or borderline estimates based on the stability of coefficient signs across resampled datasets.

Together, this framework provides a general methodological approach for robust inference of directed connectivity graph from perturbation-based experiments. This is in particular useful for high-dimensional single-cell gene expression measurements. In the following section, we recall the MRA framework and then we develop the mathematical formulation of MRA^+^ and of the network inference method; Fig. 1 provides a sketch of the two methods and the novelty in MRA^+^. In Section III, we first detail the bootstrap-based inference strategy. Then, we illustrate how the proposed framework improves the robustness and interpretability of connectivity estimates derived from perturbation-based single-cell data.

**Fig. 1.**
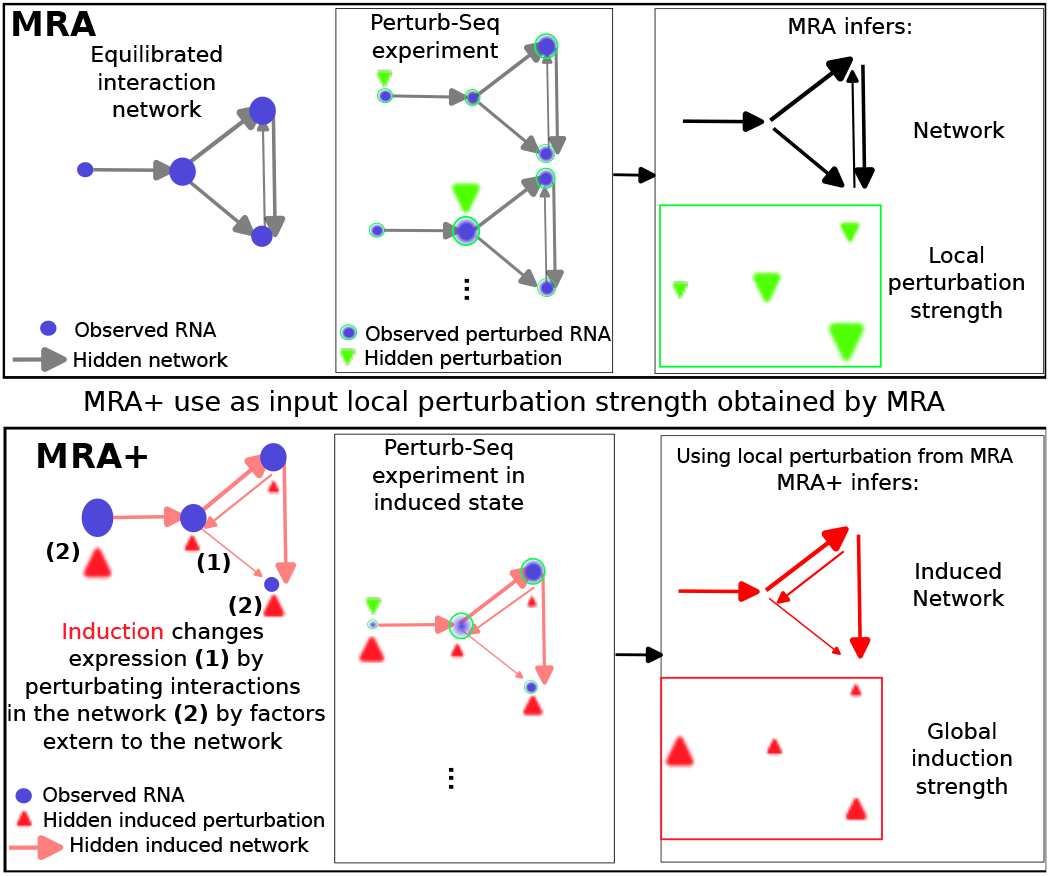
Sketches of MRA and MRA^+^. Molecular interactions between two genes may emerge or change only under specific induced conditions. **(Top)** To uncover such context-dependent connectivity shifts, the basal network is first inferred using classical MRA, where local responses to directed perturbations are estimated based on relative expression changes derived from experimental measurements. **(Bottom)** If the system is exposed to an inducing perturbation, that typically activates multiple cellular mechanisms, the hypothesis is that this global stimulus alter gene connectivity graphs and contribute to local gene expression changes through mechanisms other than direct gene-to-gene interactions. Directed perturbations are applied prior to the global inducing perturbation. This allows us to re-estimate connectivities using MRA^+^ to capture context-dependent changes. The resulting network provides both basal and induced connectivity contributions, visualized as directed edges of the graphs, thereby capturing stable as well as condition-specific regulatory edges.

## II. Extending mra to induced perturbations

### A. Classical MRA: Modular Response Analysis for gene-to-gene network inference

The fundamental definitions and mathematical aspects of Modular Response Analysis (MRA) have been established in several studies [1], [2] and then further developed in [4], [5], [9]–[11]. Given a dynamical system of *N* genes, MRA considers the linearization of this system around a steadystate. In this context, the following relationships can be written between the relative changes in gene expression 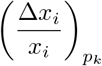 for all genes *i* when gene *k* is specifically targeted by a directed single-gene perturbation *p*_*k*_:

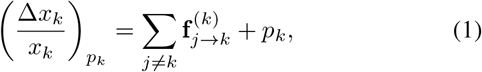

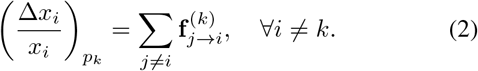

where 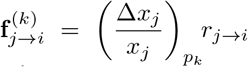 is the flux from gene *j* to gene *k*; this flux can be decomposed into the product of the relative expression change of gene *j* times the local response coefficient *r*_*j→i*_, the connectivity coefficient that captures the influence of gene *j* expression changes on gene *i*, around the steady state. In this formulation, *p*_*k*_ is the local relative expression change of gene *k* in response to its perturbation when isolated from the other genes, and it is assumed to be additive near the steady state of interest. Equation (2) tells us that the relative gene expression change for gene *i* is simply the sum of network contributions 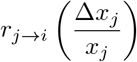 for *j* ≠ *i*, the flow of regulatory information from gene *x*_*j*_ to gene *x*_*i*_ through the underlying connectivity graph. For the perturbed gene *k*, network contributions are complemented by the local contribution *p*_*k*_ (Eq. (1)). In the context of a Gene Regulatory Network (GRN), the coefficients *r*_*j→i*_ describe the gene-to-gene regulatory connectivity of the GRN around the steady state of interest.

In general, local parameters *r*_*j→i*_ and *p*_*k*_ are not directly measurable. However, gene expression can be profiled in different experimental conditions. Noting 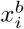 and 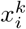 the gene expression levels in, respectively, a basal condition and in the same condition after perturbation of gene *k*, the relative changes of gene expression can be estimated from experimental data, as obtained for example using the Perturb-seq technique [8], as follows:

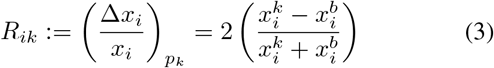

for all *i, k ∈* 1,.., *N*. This is summarized in a matrix **R** for all the experimental conditions.

Let us now define the local response matrix **r** such that *r*_*ik*_ = *r*_*k→i*_ for *i* ≠ *k* and *r*_*kk*_ = *−*1, that codes for the regulatory network. Introducing now the directed diagonal perturbation matrix **P** where *P*_*kk*_ = *p*_*k*_, the set of *N* equations (1) and (2) can be compactly written as **rR**= *−***P** which is equivalent to **r**= *−***PR**^*−***1**^, assuming **R** is invertible. Since *r*_*kk*_ = *−*1 and **P** is a diagonal matrix, identifying the diagonal terms in the latter equation leads to:

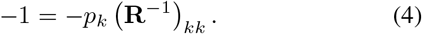

Using now the notation diag to denote the diagonal matrix *D* = diag(*M*) such that *D*_*ii*_ = *M*_*ii*_, we obtain that:

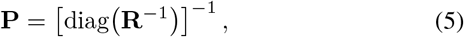

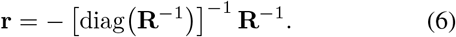

These equations are the MRA inference method, estimating the connectivity graphs **r** and the direct perturbations **P**, from the experimental data (relative gene expression changes) in **R**.

### B. The MRA^+^ model for induced perturbation

Upon exposure to a global stimulus, such as hormone stimulation or any other context-altering perturbation, the network connectivity may transition from a basal to an induced state, reflecting changes in regulatory interactions that emerge only under specific biological conditions. We hypothesize that this shift is also driven by additional local contributions triggered by the global stimulus here called 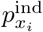, which may simultaneously influence all genes in the system. If ignored, these widespread, non-targeted effects that modulate gene expression independently of gene-to-gene interactions, would potentially bias the estimation of the underlying connectivity.

In practice, to uncover gene connectivity graphs associated to the effect of a global stimulus, genes of interests are first directly perturbed, then the cells are induced to a new biological context. To account for this, we reformulate the model of Eqs. (1) and (2) by adding the terms 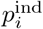 and 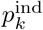 which quantify the local contributions of the induced perturbation to both directly perturbed and unperturbed genes, respectively, leading to:

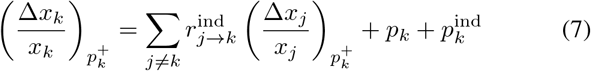

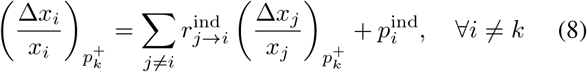

Here 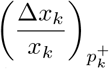 and 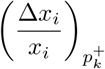 for, 1 *≤ i* ≠ *k ≤ N* denote the relative gene expression changes on *x*_*i*_ in response to a coupled perturbation: a directed perturbation on gene *x*_*k*_, combined with a multi-target inducing perturbation. This is shown by the subscript 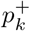.

### C. Inference in the MRA^+^ model

Since directed perturbations *p*_*k*_ are applied first, we hypothesize that their magnitude remains unchanged after the system is exposed to the global inducing stimulus. Matrix **P** can thus be determine using Eq. (5) given the relevant experimental data. This assumption also implies that the induced perturbation term is added directly to the response of the perturbed gene, as shown in Eq. (7). In contrast, unperturbed genes are affected only by the local contribution of the induced perturbation, as shown in Eq. (8). This additive structure is essential for the MRA^+^ framework, as it enables the separation of gene-specific and context-wide effects while preserving the interpretability of induced connectivity coefficients 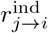 derived from combined perturbation experiments.

Let us consider the matrix **R**_ind_ that represents the relative expression changes between the directed perturbed states and the basal state, under the induced context. Each entry of **R**_ind_ captures how the expression of a gene responds to a specific directed perturbation, once the system has been exposed to the global stimulus. The matrix entries are computed from experimental data as:

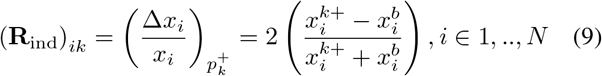

where 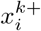 is the perturbed gene expression level of gene *i* under both the direct single-gene perturbation of gene *k* and the global (muti-target) stimuli. In the same manner as for MRA, Eqs. (7) and (8) can be written using matrix notation **r**^ind^**R**_ind_ = *−***P**_+_, where **P**_+_ *∈* ℝ^*N ×N*^. The diagonal of **P**_+_ contains the sum of directed and induced perturbations and its out-of-diagonal elements contain only induced local effects from the global stimulus. Hence, matrix **P**_+_ can be splitted into two parts:

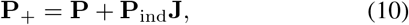

where **J** is a matrix with ones in all positions, and **P**, contains only the diagonal contributions from individually perturbed genes. For 2 genes, we would have:

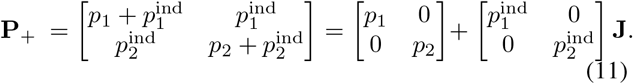

This decomposition amounts to isolate the effects of the global stimulus from the directed perturbations. The first term is the classical local perturbation matrix **P**. The second term captures the local contributions from the global stimulus, organized in the matrix **P**_ind_, which is then broadcast across all genes using the matrix **J** of ones.

This non-diagonal structure of **P**_+_ creates additional complexity in connectivity inference, as local contributions are no longer confined to directly perturbed genes. Instead, the induced context distributes regulatory input across multiple nodes in the GRN, making it necessary to disentangle network-mediated influences from inducing stimulus effects. As a result, standard diagonal inversion approaches used in classical MRA (e.g., Eq. (6)) are no longer applicable.

We thus propose a revised formulation to estimate the induced connectivity network matrix:

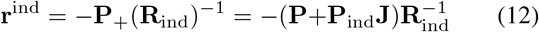

The identification of the diagonal of **r**^ind^ to *−*1 leads to the following derivation, with **I** standing for the identity:

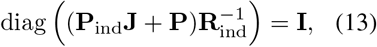

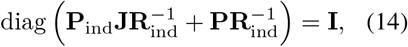

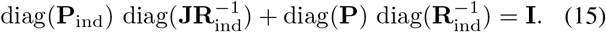

Then, because **P**_ind_ and **P** are diagonal, we have

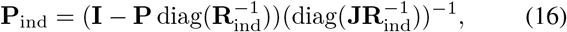

if diag 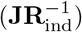 is invertible (we discuss that in III-B).

To summarise, given mean Perturb-seq type expression levels for *N* genes in a reference biological condition and after a global induction of interest, the relative expression change matrix **R** and **R**_ind_ (Eqs. (3) and (9)) can be estimated. Unless one of these matrices is ill conditioned or 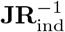 has a null diagonal term, the MRA^+^ framework then simply amounts to apply in turn equations (4), (16) and (12) in order to obtain the direct effect of the induction on each gene (**P**_ind_) and the underlying network connectivity (**r**^ind^).

## III. Application to single-cell expression data

### A. Assessing the robustness of MRA^+^ using bootstrap

The standard gene expression data, such as bulk RNA-seq data, provide the estimate of the mean expression of genes over a large cell population (typically *>* 1 million cells). Recent single-cell gene expression profiling give access to cell-to-cell gene variability but over a limited number of cells (hundreds). In this situation, mean gene expression estimation is associated with increased variability due to sampling noise and heterogeneity across cells. Therefore, it is necessary to assess both the robustness of the estimated parameters and the reliability of their interpretation.

We propose a bootstrap procedure, generating *n* bootstrap samples by sampling with replacement from the original cell population, with each sample size matching the initial number of cells. For each bootstrap sample, we recomputed the **R** matrices and re-estimated the perturbation and induced connectivity coefficients.

### B. Characterising human cells response to a virus infection

To study the behavior of the proposed MRA^+^ framework with bootstrap in a realistic setting, we analysed a gene expression dataset of primary human foreskin fibroblasts exposed to human cytomegalovirus (HCMV) infection [12]. The dataset was obtained using Perturb-seq, which combines CRISPR-based gene perturbations with single-cell transcriptomic profiling, enabling the estimation of perturbation responses across large cell populations [8]. In this context, HCMV infection acts as the inducing perturbation—introducing a complex, global stimulus that modulates the cellular regulatory landscape. We applied the MRA^+^ framework to infer the connectivity and the HCMV local influence in the infected cellular state. From the 52 genes included in the dataset, we selected the set of the 9 most expressed genes in the dataset (so as to check the method on a not too large dataset).

The inferred connectivity matrix **r**^ind^ and perturbation **P**_ind_ are on Fig. 2 A. The graph exhibited a well-defined pattern of interactions among the nine genes, in which the dominant coefficients corresponded to repressive influences on gene SEC62, counterbalancing the main direct effect of the viral infection of SEC62 activation. The median across 1000 bootstrap ensembles for the same inferred terms are in very good agreement (Fig. 2 B). This illustrates that the bootstrap samples are globally representative of the inferred coefficients.

**Fig. 2.**
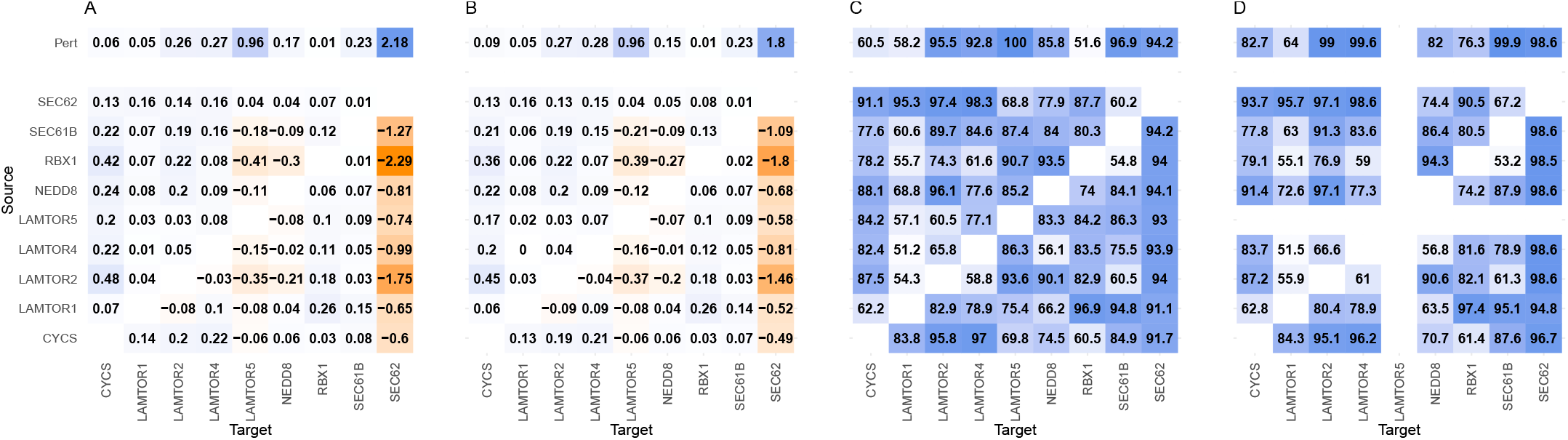
Connectivity inference and bootstrap-based robustness in the induced condition. (A) Connectivity matrix inferred with MRA^+^ for the nine-gene network, with columns representing source genes and rows representing targets; the horizontal vector indicates local induction coefficients **P**_ind_. (B) Median connectivity and induction coefficients across bootstrap resamples. (C) Percentage of bootstrap estimates whose sign agrees with the inferred coefficients, reflecting directional robustness. (D) Sign-proportion matrix after removing one gene (LAMTOR5), illustrating the effect of network composition on stability. Color intensity reflects coefficient magnitude or sign proportion; blue indicates positive values and red indicates negative values.

We applied the classic MRA framework to same dataset between the wild type cells as the basal state and the perturbation data in the infected state. The resulting connectivity matrix was significantly different than the one obtained with MRA+ (data not shown). In particular, it did not present the repression of SEC62 by the other genes in the network.

### C. Correction of an instability of the bootstrap approach

While the median is relevant, the bootstrap analysis revealed a numerical instability that we discuss now. It affects a subset of coefficients (Fig. 3). We observed that the distribution of the viral induction parameter of SEC62 is wide (Fig. 3 left column), while this is not the case for other genes, as illustrated on the typical example of LAMTOR5 (Fig. 3 right column). In particular, the inversion in the term (diag 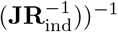 is the leading behaviour of this instability, as seen on Fig. 3, bottom left. Furthermore the fluctuations in the estimation of 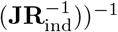 can propagate through matrix inversion and amplify variability for all terms **r**^ind^ (Eq. 16).

**Fig. 3.**
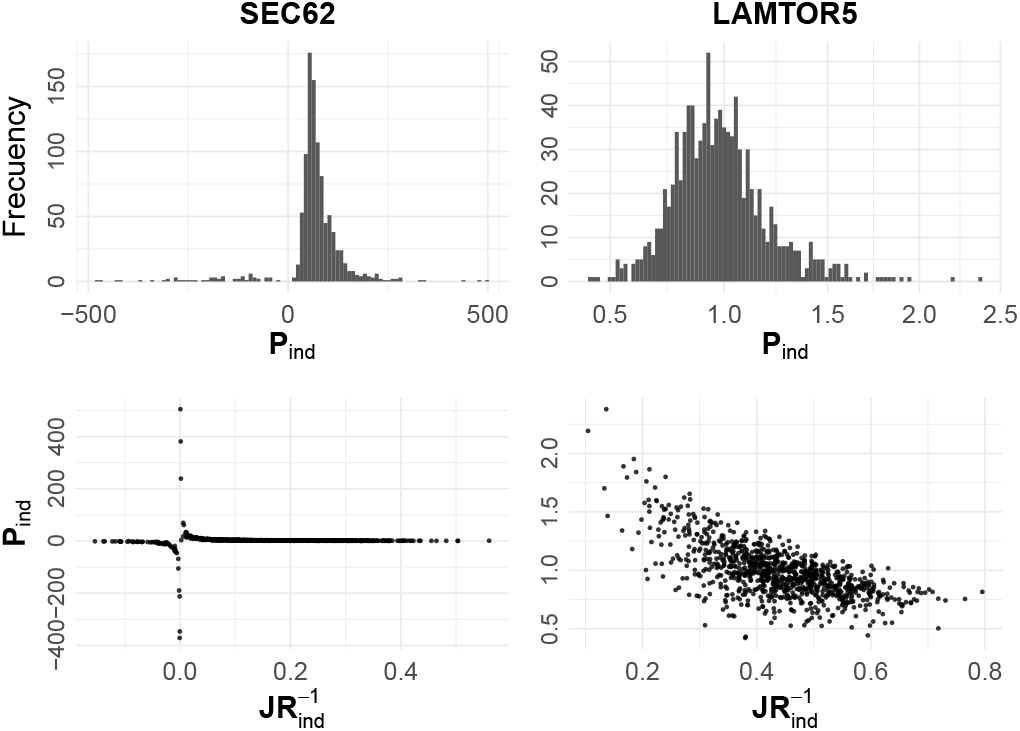
Bootstrap variability and numerical instability of induction coefficients. **Top** panels show bootstrap distributions of **P**_ind_ for two representative genes. SEC62 exhibits large variability and extreme values (mean 2, SD 29.1), whereas LAMTOR5 shows a tightly centered and stable distribution (mean 0.98, SD 0.22). **Bottom** panels display **P**_ind_ as a function of 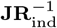. For SEC62, **P**_ind_ diverges as 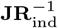 approaches zero, consistent with the inversion in Eq. (16). In contrast, LAMTOR5 remains bounded and stable. This illustrates how numerical sensitivity in intermediate terms of the MRA^+^ formulation can drive bootstrap variability and motivate complementary robustness criteria such as sign stability.

Due to the observed instabilities in **P**_ind_ estimated from Eq. (16), the standard deviation of parameters estimated from the bootstrap samples are often extremely large compared their mean value, suggesting these estimates are not usable. However, it is apparent from the distributions that nevertheless their sign was generally conserved (Fig. 3, top left). Thus, we proposed a stability analysis of coefficient signs across bootstrap samples. Specifically, we compute the proportion of bootstrap estimates whose sign is in agreement with that of the inferred coefficients **r**^ind^ from Eq. (12) computed on the sample mean. A high proportion of concordant signs indicates robust directed connectivity, even when coefficient magnitudes exhibit numerical instability. This criterion provides a practical strategy to assess the robustness and interpretability of inferred connectivity and induction coefficients.

Although several coefficients in the row of the interactions converging on SEC62 were not significant according to their standard deviation, the proportion of bootstrap samples preserving the inferred sign remained high, as seen on Fig 2 C. In particular, the connectivity coefficients directed toward SEC62, as well as the induction coefficient **P**_ind_ associated with the global perturbation, showed sign concordance exceeding 90% of bootstrap realizations. This result indicates that, despite numerical instability in magnitude, the directionality of the inferred influences remains robust, supporting the use of sign stability as a complementary criterion for interpreting borderline estimates. To validate this analysis of the signs, we removed the gene whose bootstrap estimates exhibited complete sign consistency and re-estimated the network with 8 genes, following the same (MRA^+^ and bootstrap) procedure. The network was very coherent with the previous 9 gene network (not shown) and sign stability increased substantially (see Fig. 2 D), reinforcing the confidence in the estimates.

From a biology point of view, the fact that SEC62 appears as a hub in this system is meaningful as the endoplasmic reticulum (ER)coordinates multiple stress-response pathways during viral infection. HCMV is known to remodel ER homeostasis, and SEC62 plays a role in ER stress recovery, while lysosomal signaling and proteostasis pathways are functionally coupled to ER regulation [13]–[15]. The agreement between the inferred network, local induction estimates, and bootstrap confidence with known biology indicates that MRA^+^ captures meaningful regulatory interactions in a given cellular context.

## IV. Conclusion

We introduced MRA^+^, an extension of Modular Response Analysis that incorporates global perturbations into gene connectivity inference. By explicitly modeling specific induced local contributions, the framework disentangles gene-to-gene regulatory effects from context-wide stimulus effects while preserving the interpretability of connectivity coefficients.

We showed that bootstrap resampling is essential when applying MRA^+^ to single-cell perturbation data, where sampling variability and numerical sensitivity strongly affect coefficient estimations. We identified instabilities related to the inversion of terms involving 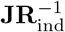 and proposed sign stability across bootstrap realizations as an effective robustness criterion. Overall, MRA^+^ provides a principled and practical framework for inferring context-dependent regulatory networks from perturbation experiments. Future work will focus on improving numerical conditioning strategies, integrating sparsity constraints and including regularisation to stabilise the analysis.

